# Marker-specific pixel attribution for single-cell protein quantification with PixAQ

**DOI:** 10.64898/2026.11.03.763929

**Authors:** Yulong Zha, Chenyu Liang, Yuxiang Lin, Guohao Li, Chen Zhang, Ziying Huang, Ze Wang, Lei Zhang, Mengsha Tong, Wenbin Jiang, Xu Xiao, Rongshan Yu

## Abstract

Imaging mass cytometry (IMC) typically quantifies all markers within the same cell mask, although their signals have distinct spatial distributions. This quantification method may omit relevant signals or include signals from neighboring cells. We present PixAQ, which improves single-cell protein quantification in IMC through marker-specific probabilistic pixel attribution. PixAQ combines spatial proximity with anchor-group intensity models to assign pixel-marker observations among neighboring cells and an explicit residual component. Original intensities are averaged over marker-specific supports. Across three cohorts (680 regions), comparisons in 366 patients showed improved agreement with predefined population-marker relationships, with cohort-median patient-paired gains of 1.99 to 4.83 percentage points in expected-marker share over shared-mask quantification. Ablation analyses showed that both spatial and intensity evidence contributed to these gains. Conventional and high-resolution IMC images linked supports to visible local and intracellular signal structures. PixAQ improves single-cell protein quantification by determining which local signals contribute to each marker measurement without changing cell segmentation.

## Introduction

Multiplexed tissue imaging measures protein expression in the spatial context of cells and tissues. Imaging mass cytometry (IMC) uses metal-tagged antibodies to map multiple protein markers in the same tissue section^1,2^. To obtain single-cell protein expression profiles, IMC studies commonly segment cells and calculate the mean intensity of each marker within the corresponding cell mask^3,4^. Together with cell coordinates, the resulting cell-by-marker matrix supports analyses of cellular phenotypes, protein co-expression and tissue neighborhoods^5–7^. Under this quantification strategy, every marker measured in a given cell uses the same set of pixels.

This shared set of pixels is determined by image-processing and segmentation procedures rather than selected separately for each marker. At cell contacts or uncertain boundaries, a mask may include signal associated with a neighboring cell or exclude signal associated with the cell being measured^8,9^. Lateral signal spillover can also occur between closely apposed cells even when their segmentation is otherwise satisfactory^8^. Because local signal distributions differ across markers, these errors can affect protein measurements differently across markers. The resulting expression profiles may still support cell phenotyping, but phenotype assignment alone does not establish whether the appropriate local signals were included in each protein measurement.

Existing approaches improve image-derived cellular measurements through mass-channel compensation^10^, improved cell segmentation^11,12^, and denoising or spatially smoothed quantification^13,14^. Other image-based methods, including Pixie, Nimbus and CellSighter, use image information for pixel-level feature annotation, marker-positivity prediction and cell classification, respectively^15–17^. Approaches addressing signal mixing between cells include REDSEA, which uses boundary-associated signals to compensate lateral spillover between adjacent cells^8^, and STARLING, which models mixtures in aggregated protein profiles to infer cellular phenotypes^18^. Related work in spatial transcriptomics has shown that molecular misassignment can distort differential expression and inferred neighborhood effects^19^, while ResolVI models local expression mixing to estimate corrected cellular expression profiles^20^. Cross-platform benchmarking of segmentation in spatial transcriptomics further shows that segmentation choices can alter transcript recovery, expression purity and downstream biological inference, including cell-type-specific expression and ligand–receptor interactions^21^. These approaches address different aspects of image analysis, cellular phenotyping and signal mixing, motivating a complementary strategy for IMC: inferring marker-specific quantification supports, defined as the sets of pixels contributing to each cellular protein measurement.

Here, we present PixAQ (Pixel Attribution for Quantification), which improves single-cell protein quantification in IMC through marker-specific attribution of local protein signals. Holding cell instances fixed, PixAQ combines spatial proximity and marker-intensity information to estimate assignment probabilities for each pixel-marker observation over nearby cells and an explicit residual component. The observation is assigned to the cell or residual component with the highest probability, allowing different markers at the same spatial coordinate to be attributed to different cells. These assignments define marker-specific quantification supports, over which original intensities are averaged to construct the cell-by-marker matrix. Across three IMC cohorts^22–24^, PixAQ improved agreement with predefined population-marker expression relationships. Analyses of a cell-line pellet of known composition^18^, conventional IMC images and selected high-resolution IMC (HR-IMC) examples^25^ linked changes in protein readouts to marker-specific supports and visible local signal structures.

## Results

### PixAQ quantifies proteins in single cells through marker-specific pixel attribution

PixAQ constructs marker-specific quantification supports for fixed cell instances through probabilistic pixel attribution. Within each region of interest (ROI), anchor groups defined by existing annotations or clustering specify which cells share a marker-specific intensity model. Throughout the analyses, Source denotes conventional shared-mask quantification, calculated as the mean raw intensity of each marker within the corresponding cell mask (Fig. 1a).

**Fig. 1.**
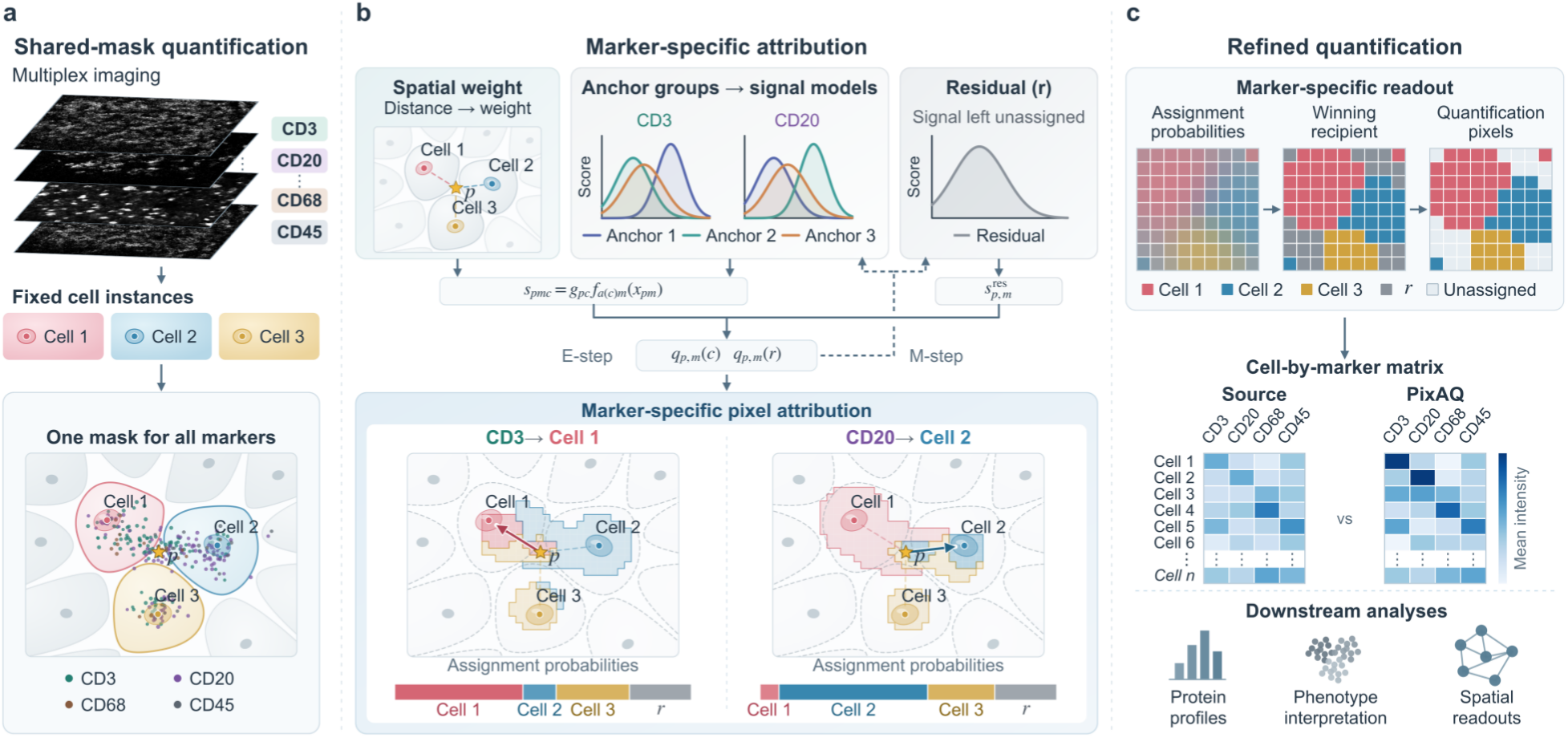
Marker-specific pixel attribution quantifies proteins for fixed cell instances. **a**, Shared-mask quantification of CD3, CD20, CD68 and CD45 (teal, purple, ochre and blue-gray dots) for fixed cell instances (Cells 1–3); every marker uses the same mask for a given cell. **b**, Marker-specific attribution with fixed cells, anchor groups and candidates. Distance-based weights and anchor-conditioned intensity scores (colored curves) define model-conditional assignment probabilities over candidate cells and residual. Lower views show CD3 and CD20 at the same coordinate (star); bars represent assignment probabilities. **c**, Hard readout: the highest-probability cell or residual determines the recipient of each pixel-marker observation. Residual winners remain unassigned. Source and PixAQ average original image intensities within shared cell masks and marker-specific supports, respectively.

For each marker intensity observed at a given pixel, hereafter referred to as a pixel-marker observation, PixAQ estimates assignment probabilities across nearby cells and an explicit residual component. Candidate cells and their spatial weights are shared across markers, whereas intensity scores depend on the marker and the anchor group of each candidate cell. PixAQ combines the spatial and intensity scores to estimate assignment probabilities and assigns each observation to the recipient with the highest probability. The residual component allows observations to remain unassigned to any cell, while marker-specific assignments allow CD3 and CD20 at the same coordinate to be attributed to different cells (Fig. 1b). For each cell-marker pair, the attributed pixels define its quantification support, and their mean raw intensity, when the support is nonempty, provides the corresponding entry in the cell-by-marker matrix (Fig. 1c). PixAQ also reports the total signal, support pixel count and whether the support is empty.

### PixAQ improves agreement with reference marker profiles

To assess whether marker-specific quantification increased agreement with the characteristic expression patterns of annotated cell populations, we defined pairs of nonoverlapping reference groups from published annotations and selected one representative marker for each group. Agreement was measured using expected-marker share. For each cell, the two marker readouts were scaled by ROI- and marker-specific factors derived from Source and held fixed across methods. Expected-marker share was calculated as the fraction of their summed scaled readouts contributed by the marker representing the cell’s reference group. Shares were averaged within each reference group, and the two group means were combined with equal weights (Methods).

We evaluated PixAQ in 680 ROIs from the Jackson breast cancer^22^, Xiao melanoma^23^ and Hoch melanoma cohorts^24^. Using Source as the baseline, we compared PixAQ with image smoothing (Blur), aggregation over transformed masks (random affine transformation intensity, RATI) and REDSEA-style correction based on cell mask boundaries. Among 366 eligible patients, median patient-paired gains in expected-marker share over Source were 1.99 percentage points in Jackson (95% patient-bootstrap interval, 1.82–2.14; 271 patients), 4.83 in Xiao (4.31– 5.60; 26 patients) and 2.67 in Hoch (2.48–3.00; 69 patients; Fig. 2a). Median changes at the patient level in expected-marker share between PixAQ and the other methods were positive in all three cohorts, with all 95% bootstrap intervals excluding zero.

**Fig. 2.**
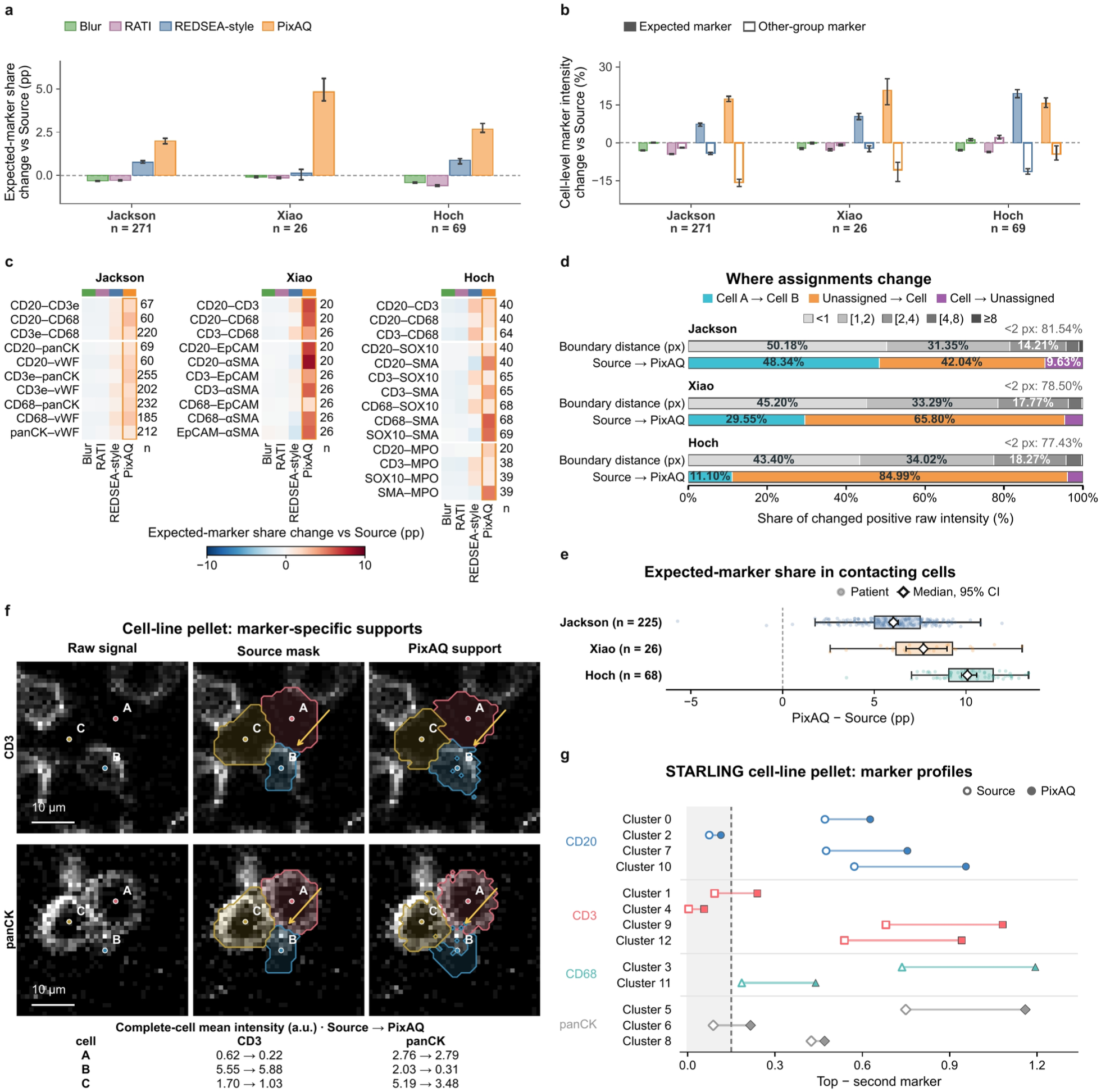
Reference-associated protein-profile changes across cohorts and in a cell-line pellet. Tissue reference labels also conditioned PixAQ. Tissue comparisons use marker-specific 95th percentiles of Source cell means within each ROI as scales, fixed across methods. **a**, Patient-median changes relative to Source in expected-marker share within predefined, Source-scaled marker pairs, averaged across reciprocal reference directions (pp); *n* = 271, 26 and 69 patients for Jackson, Xiao and Hoch. Error bars, 95% patient-bootstrap intervals. **b**, Expected-marker (filled) and other-group-marker (open) intensity changes (%); intervals as in a. **c**, Expected-marker-share changes for all 34 cohort-marker pairs, arranged by cohort with marker pairs in rows and methods in columns. Blue and red indicate decreases and increases relative to Source (pp); orange outlines mark PixAQ. Numbers at right give evaluable patient counts. **d**, Two separate marginal distributions of the same positive raw intensity involved in assignment changes, with equal ROI weights. Upper bars show distance to Source-mask boundaries (gray shades); lower bars show transition type: between-cell reassignment (cyan), assignment from outside all Source masks (orange) or loss of assignment (purple). **e**, Expected-marker-share changes in contacting cells, restricting the final PixAQ readout to each cell’s own Source mask, with marker-pair definitions as in a. Contacts are four-connected Source-mask adjacencies across opposite reference groups. Both marker readouts must be finite over nonempty intersections; measured zeros remain. Scaling and patient aggregation follow a. Points, patients; boxes, median and interquartile range; whiskers, 1.5× interquartile range; diamonds and intervals, patient median and 95% bootstrap interval. The domain includes 125,258, 152,126 and 148,922 unique cells from 225, 26 and 68 patients across 10, 10 and 14 pairs. **f**, Raw and attributed CD3/panCK signals in a 40 × 40 µm STARLING pellet patch; cells A–C were selected post hoc. Arrows mark ownership changes. The table gives paired per-cell means (a.u.). Scale bars, 10 µm. **g**, Highest minus second-highest Source-scaled mean across CD20, CD3, CD68 and panCK in 13 initial anchor clusters: Source, open; PixAQ, filled. Colors and shapes identify the Source-dominant marker: CD20, blue circles; CD3, red squares; CD68, cyan triangles; panCK, gray diamonds. Shading marks margins below a descriptive 0.15 cutoff motivated by STARLING^18^.

To examine the readout changes underlying these gains, we analyzed the expected marker and the corresponding other-group marker separately. Across the three cohorts, median changes at the patient level ranged from increases of 15.59% to 20.72% for the expected markers and decreases of 4.48% to 15.67% for the markers representing the other groups (Fig. 2b). Differences in expected-marker share between PixAQ and Source were positive for all 34 combinations of cohort and marker pair, showing a consistent direction of change across the tested marker pairs (Fig. 2c; Extended Data Fig. 1).

Assignment changes between Source and PixAQ were concentrated near the cell mask boundaries across all ROIs. In Jackson, Xiao and Hoch, respectively, 81.54%, 78.50% and 77.43% of the changed intensity was located within two pixels from the cell mask (Fig. 2d). We then classified the assignment changes by their transition from Source to PixAQ, including reassignment between cells, assignment of signals from outside all Source masks to a cell, and loss of an existing cellular assignment. The composition of these changes differed across cohorts.

In Jackson, reassignment between cells accounted for 48.34%, while signals assigned from outside cell masks accounted for 42.04%. The latter category was more prominent in Xiao and Hoch, accounting for 65.80% and 84.99% of the changed intensity, respectively.

The substantial contribution of signal from outside Source masks raised the question of whether the improvement in expected-marker share depended on signals outside the masks. We therefore focused on reference cells contacting cells from the paired reference group and recalculated PixAQ means using only the intersection of each cell’s marker-specific support with its own Source mask, without refitting the model or changing pixel assignments (Methods). Within this restricted cell set, expected-marker share remained higher than Source for the same cells by 6.04, 7.68 and 10.08 percentage points in Jackson, Xiao and Hoch, respectively (Fig. 2e; Supplementary Table 1). These results show that PixAQ improved agreement with expected marker patterns even when quantification was restricted to signals within the original mask.

The STARLING cell-line mixture pellet^18^ provided an additional setting in which to examine marker-specific changes in a sample of known composition. The sample contained 2,712 cells, with its 13 initial clusters used as PixAQ anchors. In a selected region, cells showed different assignment changes for CD3 and panCK (Fig. 2f). For cell B, PixAQ included more CD3 signal around the cell, making its local CD3 pattern more distinct and increasing its mean intensity from 5.55 to 5.88. By contrast, PixAQ excluded some panCK signals to neighboring cells A and C, reducing its mean intensity from 2.03 to 0.31. For cell C, PixAQ excluded peripheral pixels in both channels while retaining those in the central region. This example illustrates the potential for peripheral signal loss when its cellular attribution is uncertain.

We then extended the analysis to protein profiles across all 13 initial clusters. We defined the margin as the difference between the highest and second-highest scaled mean intensities among CD20, CD3, CD68 and panCK (Methods). PixAQ increased this margin by 0.243 on average, and the number of clusters with a margin below 0.15 decreased from four to two. No cluster changed its dominant marker (Fig. 2g; Supplementary Table 4). PixAQ therefore strengthened dominant-marker contrast while preserving the existing dominant-marker patterns.

### Spatial and intensity evidence jointly determine marker-specific pixel attribution

We next examined how spatial weighting, marker-intensity information and anchor-group correspondence contributed to the observed gains in reference-profile agreement. Spatial weights favor nearby candidate cells, whereas marker-intensity scores reflect the compatibility of each observation with the candidate cell’s anchor-group intensity distribution. At a single coordinate in a melanoma ROI, CD20, CD4 and CD127 shared the same candidate cells and spatial weights, yet their marker-specific intensity scores yielded different highest-probability recipients: C2, C1 and the residual component, respectively (Fig. 3a,b).

**Fig. 3.**
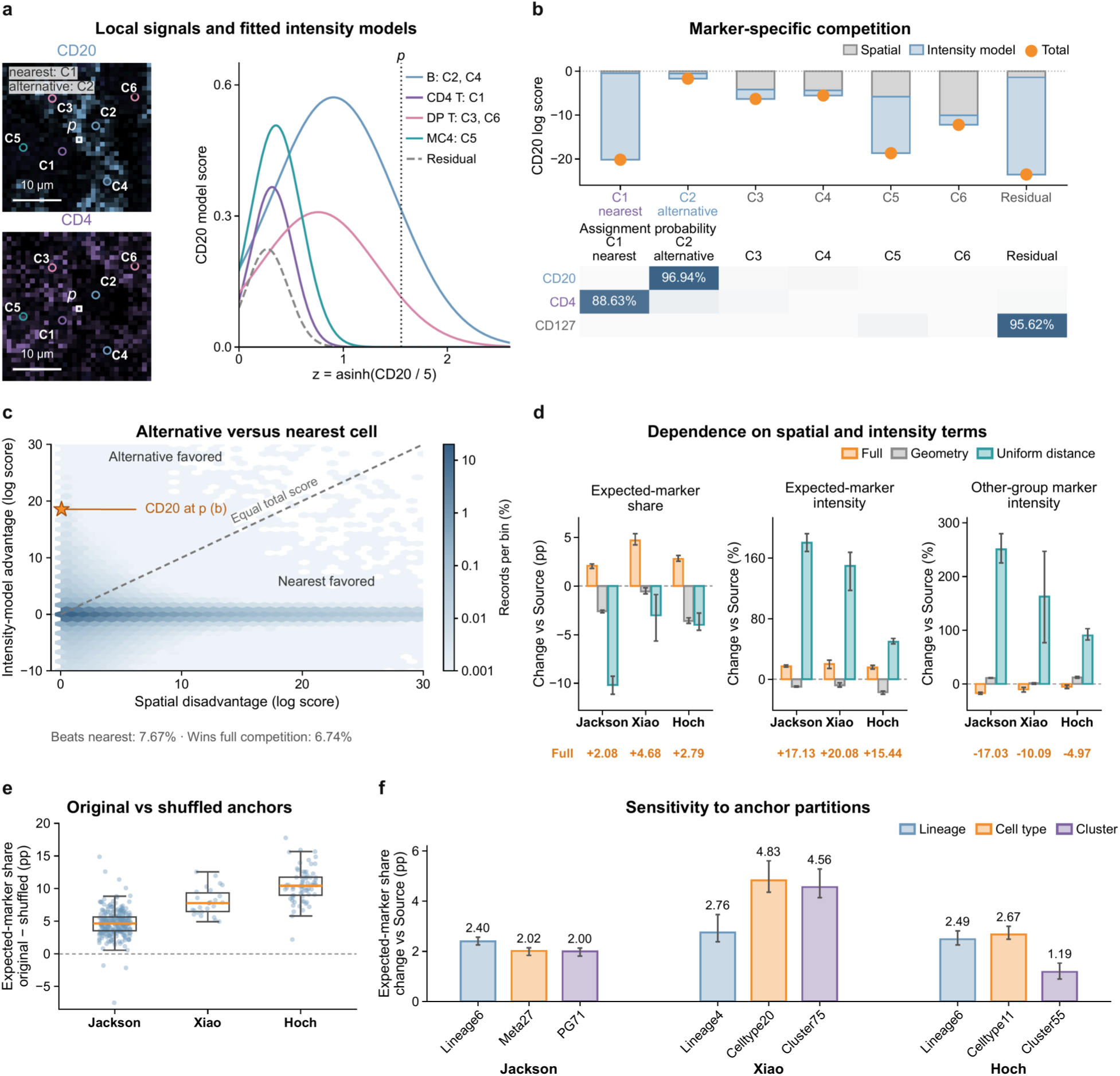
Marker-specific competition, component dependence and anchor-partition sensitivity. **a**, Raw CD20/CD4 and dim DNA around pixel p and six candidates, with fitted anchor-group and residual CD20 scores *ρφ*(*z*; *μ*, *σ*), where *z* = asinh(*x*/5) for positive intensity. Curves are not renormalized; the vertical line marks *z*(*p*) = 1.56. Scale bars, 10 µm. **b**, Spatial, intensity-model and total log scores for CD20, and three-marker assignment probabilities; residual winners remain unassigned. **c**, Intensity-model advantage (alternative minus nearest) versus spatial disadvantage (nearest minus alternative), in log-score units. The alternative maximizes the combined spatial-intensity log score among candidates outside the nearest cell’s anchor group. Dashed line, equal total scores; star, CD20 in b. Hexagon colors and frequencies use 3,928,035 positive observations with an alternative from a different anchor group, across six markers and 137 Xiao ROIs, sampling 20,000 positions per ROI with equal weights across 822 ROI-marker strata. The alternative beats the nearest cell in 7.67% and wins against all candidates and residual in 6.74%. **d**, Expected-marker-share (pp) and expected-/other-group-marker intensity changes (%) relative to Source. Full uses fine anchors; Geometry removes candidate-cell intensity scores; Uniform distance equalizes spatial weights while preserving their total. Arms were fitted separately with fixed candidates and residual competition. Common eligibility across these arms, Source, Lineage and Fine shuffled (within-ROI fine-label permutation) yields *n* = 257, 26 and 69 patients in Jackson, Xiao and Hoch. Intensity contrasts are back-transformed from log_2_. **e**, Patient-paired fine-original minus fine-shuffled share changes; patients as in d. Points, patients; boxes, median and interquartile range; whiskers, 1.5× interquartile range. **f**, Lineage, annotated cell-type and published cluster anchors under matched fitting settings and common eligibility; *n* = 271, 26 and 69 patients. Only anchor memberships varied (Supplementary Table 3). Bars in d/f, cohort medians; intervals, 95% patient-bootstrap intervals.

This example illustrates how marker-specific intensity evidence can change attribution under the same spatial configuration. We then examined how often a candidate from a different anchor group could overcome the spatial preference for the nearest cell. Across 137 Xiao ROIs, we uniformly sampled 20,000 pixel positions per ROI and evaluated six markers, retaining 3,928,035 positive-signal pixel-marker observations with at least one candidate from an anchor group different from that of the nearest cell. For each observation, we selected the highest-scoring alternative from these groups using the combined spatial-intensity log score. This alternative outscored the nearest cell in 7.67% of cases and won against all candidates and the residual component in 6.74%, becoming the final recipient. Both rates were averaged equally across 822 ROI-marker strata (Fig. 3c). These results demonstrate that marker-intensity information can overcome distance-based preference and alter the final recipient of local protein signals.

We next tested whether spatial weighting and marker-intensity information both contributed to the observed gains in reference-profile agreement. We compared the full model with two separately refitted component controls. Geometry removed candidate-cell intensity scores while retaining distance-dependent spatial weights. Uniform distance equalized spatial weights among candidate cells at each pixel while preserving their total. Both controls retained the original candidate sets and competition with the residual component. Among cells eligible across all controls, expected-marker share was lower than Source by 0.55–3.58 percentage points with Geometry and 3.01–10.17 percentage points with Uniform distance (Fig. 3d; Extended Data Fig. 3). Although Uniform distance increased mean intensities of both expected and other-group markers by 49.69–250.61%, it did not reproduce the full model’s gains in expected-marker share. These results indicate that both distance-dependent spatial weighting and candidate-cell intensity evidence contributed to the observed improvement in reference-profile agreement.

Because intensity scoring uses anchor-group expression patterns, we next examined the contribution of cell-to-group correspondence. We randomly shuffled fine anchor labels within each ROI while preserving group sizes. Expected-marker share then fell below Source in all three cohorts; median patient-paired original-versus-shuffled differences were 4.66, 7.79 and 10.44 percentage points in Jackson, Xiao and Hoch, respectively (Fig. 3e). Shuffled groups did not reproduce the gains in reference-profile agreement, indicating that retaining the original cell-to-anchor-group correspondence contributed to the observed improvement.

Beyond cell-to-group correspondence, we compared anchor granularity under matched fitting settings, using lineage groups, annotated cell types and published clusters. All three levels improved expected-marker share over Source, but gains did not increase monotonically as groups became finer (Fig. 3f; Supplementary Table 3). In Xiao, gains were 2.76, 4.83 and 4.56 percentage points for Lineage4, Celltype20 and Cluster75, respectively. Jackson and Hoch likewise showed no consistent pattern of larger gains with finer groups.

### Marker-specific pixel attribution links local protein signals to single-cell quantification

Having examined how spatial and intensity evidence affected assignments, we next compared PixAQ quantification supports with visible protein-signal structures in Xiao melanoma. In a selected tissue field, raw signals, Source masks and PixAQ supports showed that some locally visible signal structures extending beyond Source masks were incorporated into PixAQ supports (Fig. 4a). We then visualized seven marker channels illustrating reassignment between cells, new inclusion in cellular supports and loss of assignment relative to Source (Fig. 4b). Among 4,149 coordinates with positive raw signals for both CD20 and CD4, 953 (22.97%) had the two markers assigned to different cells (Extended Data Fig. 4), illustrating marker-specific attribution at the same spatial coordinate.

**Fig. 4.**
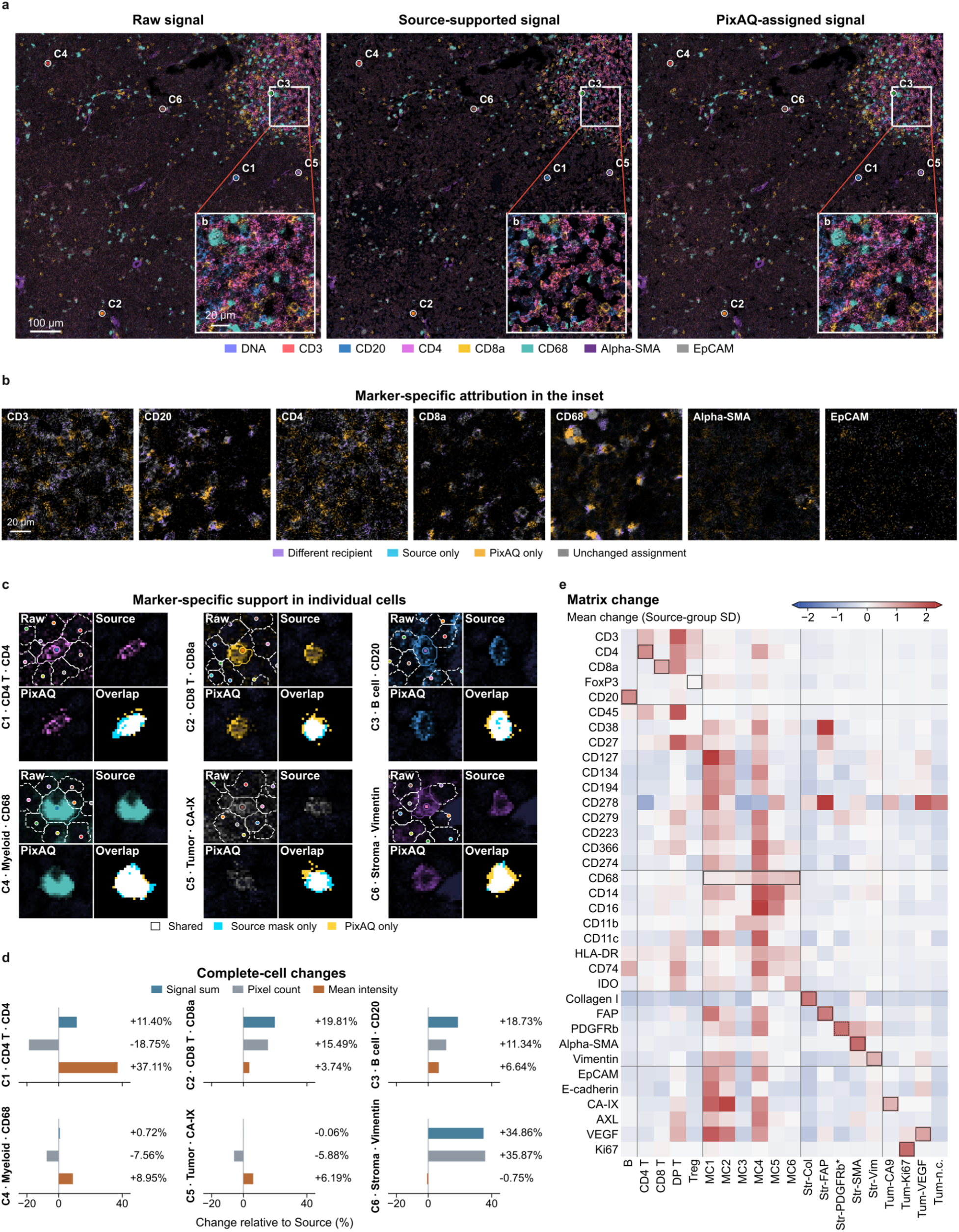
Linking marker-specific assignments to cellular protein profiles in melanoma. All panels use Celltype20 anchors. **a**, Raw, Source-supported and PixAQ-assigned signals in ROI 64F_ROI10; c1–c6 locate the cells in c. **b**, Seven inset marker channels showing assignment relations: purple, different recipients; cyan, Source only; amber, PixAQ only; gray, identical recipients or unassigned in both methods. Brightness represents raw intensity. **c**, Six fixed cell-marker examples showing raw signal, Source/PixAQ supports and overlap in 32 × 32 native-pixel windows. Colored solid outlines mark target Source masks; white dashed outlines mark neighbors; hollow circles mark cell centers. **d**, Percentage changes relative to Source in per-cell signal sum (A, blue), support pixel count (N, gray) and mean intensity (Q = A/N, orange) for c1–c6, over each method’s full quantification support, including pixels outside the crops. Each case is one cell-marker measurement. **e**, ROI-balanced reference-group mean changes, scaled by the standard deviation across Source group means; zero denotes no change. Outlines mark reference-associated entries; colors saturate at ±2.3. The asterisk and dashed outline flag PDGFRβ-positive stroma, represented by eight eligible ROIs. Scale bars: a, 100 and 20 µm; b, 20 µm.

We further examined six cell-marker examples across T-cell, B-cell, myeloid, tumor and stromal populations, linking local raw images and quantification supports (Fig. 4c) to per-cell signal sums, support pixel counts and mean intensities (Fig. 4d). For CD8a, CD20 and Vimentin, PixAQ supports followed visible signal structures, including portions extending beyond Source masks. For Vimentin, signal sum and support pixel count increased nearly proportionally, by 34.86% and 35.87%, leaving the mean almost unchanged (−0.75%).

For CD4, PixAQ retained the bright central signal structure and included 11.40% more raw signal than Source despite using 18.75% fewer pixels, increasing mean intensity by 37.11%. CA-IX and CD68 showed more localized adjustments to pixel inclusion. Their signal sums remained nearly constant (−0.06% and +0.72%, respectively), while support pixel counts decreased by 5.88% and 7.56% and mean intensities increased by 6.19% and 8.95%. These contrasting cases illustrate marker-specific support selection without changing the original cell instances. Mean-intensity changes alone cannot establish attribution plausibility; the selected signal structures, signal sums and support sizes must also be considered. Similar comparisons in Jackson and Hoch linked local quantification supports to per-cell signal sums, pixel counts and mean intensities (Extended Data Fig. 5). All measurements were calculated over each method’s complete quantification support, including pixels outside the displayed image crops.

Moving from individual-cell examples to population-level profiles, we examined how PixAQ altered protein readouts across the 20 Xiao reference groups. For each of 35 markers, we calculated ROI-balanced PixAQ-minus-Source differences in group-mean intensity, normalized by the standard deviation of Source group means for that marker (Fig. 4e). Changes varied in direction across the 700 marker-group combinations, with lower PixAQ readouts in 459. Mean intensities increased for all 12 prespecified marker-group associations, including CD20 in B cells and CD4 in CD4 T cells. However, increases were not restricted to these associations: some myeloid groups showed higher mean readouts for both lineage-associated and tumor-associated markers. These patterns indicate that PixAQ produced marker- and population-dependent changes rather than a uniform shift in protein intensities.

We further assessed whether these multi-marker readout changes affected discrimination among five published reference classes: T cells, B cells, myeloid cells, stromal cells and tumor cells. In an auxiliary patient-held-out classifier analysis, macro-F1 increased from 0.566 for Source to 0.713 with Cluster75 anchors (paired gain, 0.1465; 95% interval, 0.1331–0.1582) and reached 0.737 with Celltype20 anchors (Extended Data Fig. 6a,b). PixAQ was fitted to the full cohort before classifier evaluation, with patient-held-out splits applied only to the downstream classifier. Inspection of selected clusters also revealed heterogeneous changes in marker readouts (Extended Data Fig. 6c–h). Together, these analyses show that marker-specific support selection changes both individual-cell readouts and population-level protein profiles, with effects varying across markers and cell populations.

### Marker-specific supports correspond to visible protein-signal structures in HR-IMC

Although conventional IMC allowed us to examine local attribution changes, finer protein-signal structures were less clearly resolved. We therefore extended the analysis to high-resolution IMC (HR-IMC), which provides a more detailed view of local and intracellular protein-signal distributions^25^. We applied PixAQ to 4,756 fixed cell instances across three HR-IMC regions, using clustering-derived anchor groups and shape-aware spatial weighting for eligible cells (Methods). In ROI001, we compared raw protein signals with the portions included by Source masks and PixAQ supports to examine how the two quantification approaches captured visible structures (Fig. 5a).

**Fig. 5.**
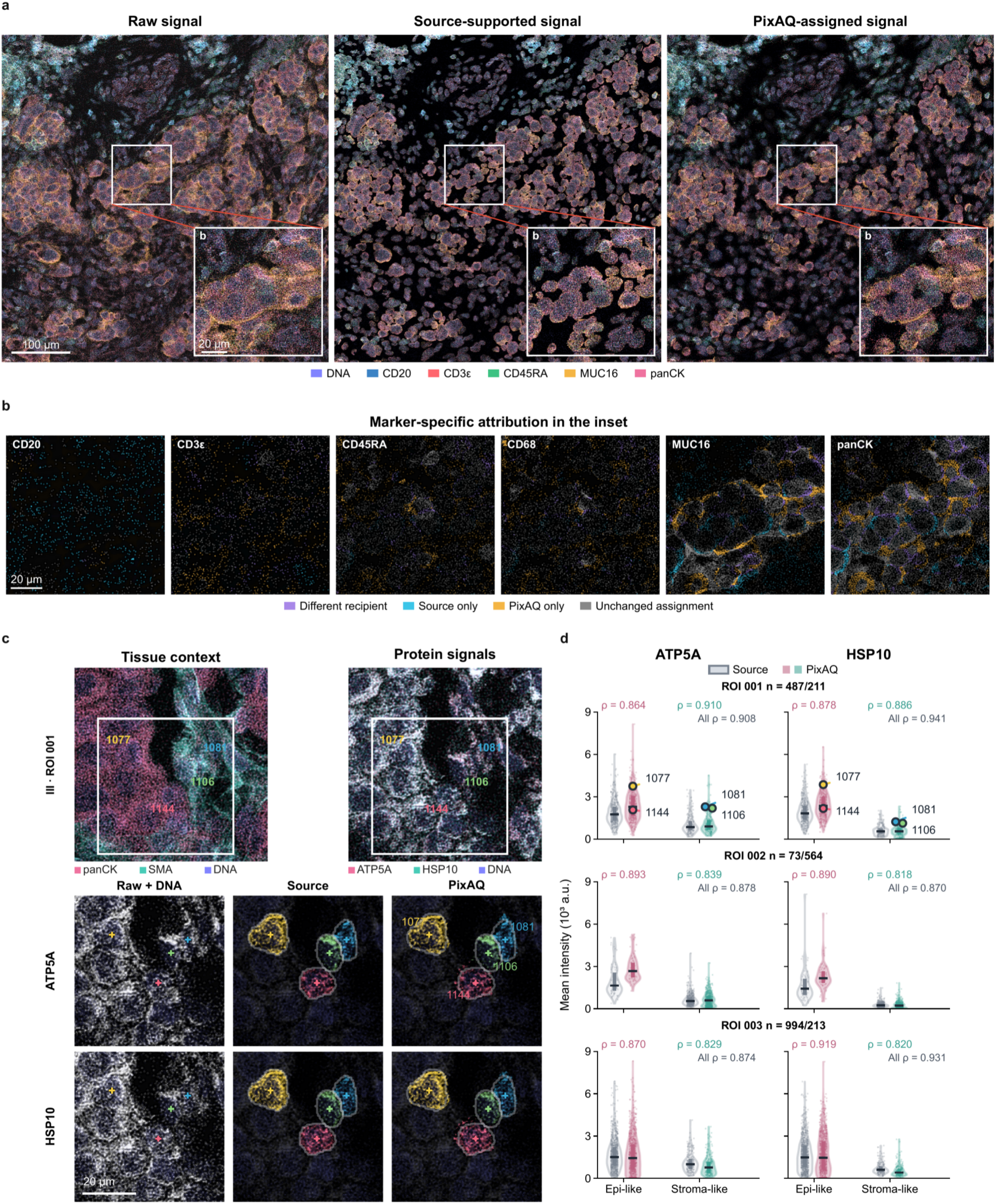
Marker-specific pixel attribution and protein quantification in HR-IMC. All panels use fixed cell instances and the PG23 shape-aware fit; Source masks provide the structural and pooling comparator. **a**, Raw signals, signals within any Source mask and signals assigned to any cell by PixAQ in ROI001, with the boxed patch enlarged at the lower right. Marker colors and display scales are shared across views; DNA provides unchanged spatial context. **b**, Six inset marker channels showing assignment relations: purple, different recipients; cyan, Source only; amber, PixAQ only; gray, identical recipients or unassigned in both methods. Brightness represents raw intensity. **c**, Field III (ROI001): ATP5A/HSP10 patterns in panCK- and SMA-associated tissue context, with raw signal and Source/PixAQ supports for four numbered cells. Support colors identify cells; gray, raw context; purple, DNA; pale outlines, Source masks. **d**, ATP5A/HSP10 mean raw intensities (10³ a.u.) over each method’s full quantification support for 2,542 cells with measured Source-mask areas of 500–8,000 pixels. Epithelial-like and stromal-like groups were defined from PG23 Source protein profiles. Group sizes are 487/211, 73/564 and 994/213 in ROIs 001–003, respectively. Source is gray; PixAQ is colored. Violins show distributions; points, individual cells; ticks, medians; bars, interquartile ranges. Spearman ρ compares paired Source/PixAQ readouts within each ROI and marker, across all displayed cells (All ρ) and within each group (colored ρ). Numbered points correspond to c. Scale bars: 100 µm (a overview) and 20 µm (a inset; b,c).

In the enlarged ROI001 field, Source masks excluded portions of a visible MUC16 arc and adjacent panCK structures, whereas PixAQ supports incorporated more of these signals into cellular measurements (Fig. 5a). The corresponding six-marker maps further distinguished signals reassigned between cells from those included in cellular measurements by only one method (Fig. 5b). A complementary immune-contact example showed concentrated CD20 signal around cell 1423 extending toward neighboring cells 1388 and 1446. Relative to Source, PixAQ increased the CD20 signal sum attributed to cell 1423 by 27.13%, while reducing the corresponding sums in the two neighboring cells by 98.84% and 95.32%, respectively (Extended Data Fig. 7d). The accompanying four-marker readouts illustrated how these changes in local signal attribution were reflected in the respective cellular protein measurements (Extended Data Fig. 7e).

Having examined marker-specific attribution at cell contacts, we turned to intracellular protein-signal patterns. We focused on ATP5A and HSP10, which were used to map mitochondrial networks in the original HR-IMC study^25^. In four numbered cells from ROI001, we compared raw marker signals, Source masks and PixAQ supports, with panCK, SMA and DNA providing tissue and cellular context (Fig. 5c). The marker-specific supports corresponded to visible intracellular signal patterns, with Source-mask outlines shown for comparison. Similar spatial correspondence was observed in additional cell examples from two other HR-IMC regions, alongside their corresponding readouts over each method’s full quantification support (Extended Data Fig. 7a–c).

Beyond these local examples, we assessed how the revised quantification supports related to cellular protein readouts across a larger set of cells. We compared PixAQ and Source mean intensities for ATP5A and HSP10 in 2,542 cells across three HR-IMC regions (Fig. 5d). These cells met the Source-mask area criterion and belonged to two descriptive groups, epithelial-like and stromal-like, defined from existing Source protein profiles (Methods). The numbered cells from Fig. 5c were highlighted in the corresponding distributions. PixAQ and Source readouts showed strong cell-wise rank correlations for both markers in each ROI (Spearman’s ρ = 0.870–0.941), which remained strong within each descriptive group (ρ = 0.818–0.919). In all three regions, both methods yielded higher median intensities for ATP5A and HSP10 in epithelial-like than in stromal-like cells.

Together, across three IMC cohorts, PixAQ improved agreement with predefined population-marker expression relationships, while conventional and high-resolution IMC image analyses linked marker-specific quantification supports to visible local and intracellular protein-signal structures. These findings support improved single-cell protein quantification through marker-specific pixel attribution, allowing local spatial and marker-intensity information to determine which signals contribute to each measurement without redefining the cells themselves.

## Discussion

PixAQ improves single-cell protein quantification in IMC through marker-specific probabilistic pixel attribution. Across three IMC cohorts, quantified profiles showed closer agreement with predefined population-marker expression relationships. In a cell-line pellet of known composition, dominant-marker contrasts increased on average within existing clusters without changes in their dominant markers, while image analyses across multiple cohorts linked marker-specific supports to visible local protein-signal structures. Together, these findings highlight a measurement problem distinct from cell identification and phenotyping: a protein profile sufficient for broad lineage assignment does not, by itself, establish that the local signals contributing to each protein measurement have been appropriately selected.

PixAQ addresses this problem by retaining existing cell instances while inferring marker-specific quantification supports. Anchor-group intensity distributions provide a reference for assessing the compatibility of local observations with candidate cells, while spatial weights distinguish nearby cell instances. Cells sharing an intensity model can therefore receive different observations according to their spatial relationships, and an explicit residual component allows observations to remain unassigned. Assignment probabilities are converted into hard supports, over which original intensities are averaged. Anchor-group information thus influences cellular protein measurements through signal selection, rather than by replacing individual readouts with group-level expression estimates. This allows local spatial and marker-intensity information to participate directly in constructing each cellular protein measurement.

Because PixAQ changes which observations enter each measurement, the resulting mean intensity depends on both the included signal and the size of its quantification support. Uniform distance increased mean intensities without reproducing the full model’s gains in expected-marker share, whereas the Vimentin example incorporated more visible signal over a larger support with little change in mean intensity. These observations show that the direction or magnitude of a mean change alone cannot establish attribution plausibility. Signal sums and support pixel counts help explain how each mean is constructed, while conventional and high-resolution IMC images provide spatial evidence relating the selected supports to visible local and intracellular protein-signal structures. The resulting readouts represent mean original intensities over inferred marker-specific supports, rather than calibrated estimates of whole-cell protein abundance.

This support-based approach nevertheless has limitations arising from the fixed cell instances and the attribution model. PixAQ cannot recover cells missing from the supplied cell instances, and inaccuracies in cell centers can affect candidate selection and attribution. Each pixel-marker observation is assigned to a single cell or the residual component, without separately estimating contributions from multiple cells within that observation. Although PixAQ estimates assignment probabilities, the final hard-support readout does not propagate this uncertainty into the cell-by-marker matrix. Residual observations may also contain biological signal. Comparisons with raw protein images provide evidence for the spatial correspondence of marker-specific supports with visible structures but do not independently establish the cellular origins of individual pixel-marker observations. Direct validation of pixel-level attribution accuracy will require independent reference measurements.

A further consideration is the dependence of signal attribution on anchor-group information. PixAQ does not require biologically annotated cell types, as precomputed clusters can also define anchor memberships. However, incorrect group assignments or poorly represented rare expression states may bias attribution. Shuffling cell-group correspondence did not reproduce the gains in reference-profile agreement, whereas finer anchor partitions did not consistently yield larger improvements. These findings indicate that informative cell-group correspondence matters for attribution, while finer grouping alone does not guarantee better results. This dependence also affects evaluation: in the annotated tissue cohorts, anchor construction and reference assessments drew on the same published annotations. Greater agreement with predefined population-marker relationships therefore supports reference-profile consistency but does not independently establish quantitative accuracy. Similarly, increased separability of reference classes does not by itself demonstrate improved phenotyping. Because PixAQ was fitted to the full cohort before patient-held-out classifier evaluation, these results do not establish generalization of the complete workflow to unseen patients.

Reducing reliance on predefined anchor memberships is therefore a natural direction for further development. One possibility is to jointly infer cellular expression states and marker-specific pixel attribution. ResolVI^20^ provides a related precedent at the aggregated-expression level by jointly modeling latent cellular states and local signal mixing in spatial transcriptomics. Extending this principle to pixel-resolved IMC could allow inferred cell states and signal assignments to inform each other while retaining existing cell instances. Such joint inference would require safeguards against mutually reinforcing errors between states and assignments, together with independent validation of quantitative accuracy and preservation of genuine cellular heterogeneity. The present study demonstrates that single-cell protein quantification can be improved by allowing marker-specific spatial and intensity evidence to determine which local signals enter each measurement, without redefining the cells themselves.

## Methods

### Datasets and model inputs

PixAQ takes multichannel protein images, cell instance identifiers and center coordinates, and fixed anchor-group memberships as inputs. Anchor groups organize the estimation of marker-intensity distributions and can be supplied by annotated populations or precomputed clusters. Cell instances, centers and group memberships remain fixed during fitting. Given these fixed inputs, Source-mask boundaries are not used in candidate generation, intensity-model fitting or pixel attribution. The masks define the Source quantification comparator and are also used in the mask-based evaluation analyses described below.

We analyzed 376 Jackson breast-cancer regions of interest (ROIs)^22,26^, 137 Xiao melanoma ROIs^23,27^, 167 Hoch melanoma ROIs^24,28,29^, one STARLING cell-line pellet containing 2,712 cells^18,30^, and three Bollhagen high-resolution IMC (HR-IMC) ROIs containing 4,756 cells^25,31^. Published images and cell instances were retained, and images were analyzed at native resolution. Throughout, raw intensity denotes the supplied image intensity before PixAQ processing, including the published spillover correction for Hoch.

Tissue analyses used Jackson Lineage6/Meta27, Xiao Lineage4/Celltype20 and Hoch Lineage6/Celltype11 as lineage and fine anchor partitions. Published cluster partitions—Jackson PG71, Xiao Cluster75 and Hoch Cluster55—were reused without reclustering. These partitions remained related to the published annotations used for evaluation. Their origins, marker inputs and coverage are documented in Supplementary Table 3.

The pellet used STARLING’s 13 initial PhenoGraph^32^ clusters, which preceded its subsequent cell-type assignments^18^. HR-IMC anchors were reconstructed using the published seven-marker PhenoGraph workflow^25^. Clustering the unscaled mean intensities of Vimentin, CD45RA, pan-cytokeratin, SMA, β-catenin, E-/P-cadherin and CD31 in 3,367 cells with deposited areas of 500–8,000 pixels yielded 23 groups (PG23; *k* = 20). The remaining 1,389 cells were assigned by majority vote among their 20 nearest reference cells in the same expression space. Anchor construction used no PixAQ measurements.

### Marker-specific pixel attribution

For pixel *p*, the candidate set C(*p*) contains up to six nearest cell centers within a specified radius and is shared across markers. The spatial weight for candidate cell *c*, centered at ***μ****_c_*, is

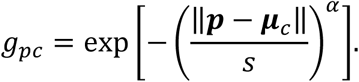

Tissue IMC and pellet analyses used a candidate radius of 42 pixels, *s* = 6 pixels and *α* = 2.5. HR-IMC used a radius of 126 pixels and *s* = 16 pixels at 0.333 µm per pixel, with the same exponent.

Within each ROI, cells in the same anchor group share a marker-intensity model. For raw intensity *x*, positive values are transformed as *y_m_*(*x*) = asinh(*x*/*k_m_*), with *k_m_* = 5 for tissue IMC and HR-IMC and *k_m_* = 1 for the pellet. Suppressing the ROI index, the intensity score for anchor group *t* and marker *m* is

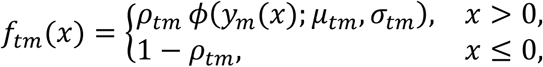

where *ρ_tm_* is the detection probability and *φ* is a Gaussian density. For positive observations, this density is used as a working score on the transformed scale without renormalization to the positive half-line. A separate ROI- and marker-specific residual model, *f*_res,*m*_, has the same form.

Let *a*(*c*) denote cell *c*’s anchor group. Candidate-cell and residual assignment probabilities are

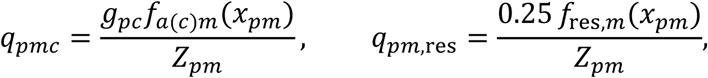

with

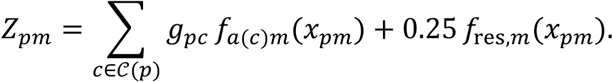

Competition is evaluated separately for each marker. Candidates sharing an anchor group have the same intensity score, so their relative preference depends on spatial weights. Candidates from different groups can also differ in intensity score. These probabilities represent model-conditional relative support among candidate cells and residual.

### Parameter estimation and fitting-budget assessment

Intensity models were fitted separately for each dataset using an EM-style procedure^33^, alternating assignment calculations with weighted updates of detection probabilities, positive-intensity means and variances. For each observation, an anchor group’s weight was the sum of assignment probabilities over candidates belonging to that group; the residual weight was *q_pm_*_,res_. Anchor models were initialized from four-pixel-radius neighborhoods around cell centers. Residual initialization used the lower-detection component of a two-component mixture fitted at pooled and ROI levels.

Detection probabilities were normalized by total assignment weight, whereas positive-intensity moments were normalized by weight on positive observations. ROI-specific anchor estimates with zero relevant weight fell back to their pooled estimates. For pooled anchor estimates and residual estimates at both ROI and pooled levels, including residual initialization, zero total assignment weight yielded a detection-probability estimate of zero before clipping. Zero assignment weight on positive observations yielded a positive-intensity mean estimate of zero and a variance equal to the squared standard-deviation floor before any ROI-level shrinkage.

ROI parameters were shrunk toward pooled estimates:

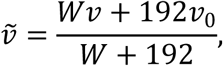

where *v* and *v*_0_ are the ROI and pooled estimates. For anchor models, *W* is the group’s total posterior weight within the ROI; for residual models, it is the ROI pixel count. Variances were shrunk before taking square roots. Detection probabilities were clipped to [10^−4^,1 − 10^−4^], and standard deviations were floored at 0.2.

Primary analyses used ten M-step updates for Jackson and Xiao, twenty for Hoch, and five for the pellet and HR-IMC, followed by a final E-step for attribution. We monitored the sum of log score-normalizers, ∑*_p_*_,*m*_ log *Z_pm_*, without a shrinkage penalty. The procedure is described as EM-style because the regularized moment updates do not exactly maximize this objective.

Fitting-budget sensitivity was assessed with matched short and extended fits across all tissue ROIs (Extended Data Fig. 8). Within each cohort, images, cells, fine-anchor memberships, implementation, initialization, spatial settings and seed were held fixed. Budgets were 10 versus 100 updates for Jackson and Xiao and 20 versus 100 for Hoch. Extended fits permitted early stopping after at least eight updates when the absolute relative objective change and maximum relative parameter change were at most 10^−4^ and 10^−3^, respectively, for three consecutive updates. Parameter change was measured by the relative *L*_2_ change of each ROI-level parameter block and maximized across active anchor and residual blocks.

### Quantification supports and cellular readouts

Each pixel-marker observation is assigned to the candidate cell or residual component with the highest final probability. Cell-residual ties favor the cell; residual winners and observations without candidates remain unassigned. No connected-component filtering is applied after attribution.

The finite-intensity support for cell *c* and marker *m* is

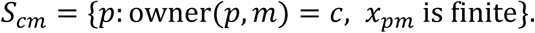

Signal sum, support pixel count and mean intensity are

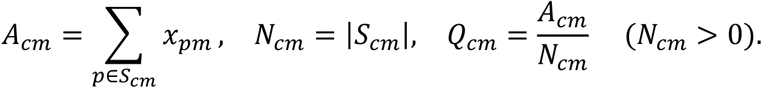

Source applies the same calculations within the supplied cell masks. PixAQ matrix entries are unweighted means of original intensities over marker-specific supports. Assigned measured zeros contribute to *N_cm_*, so a nonempty all-zero support has a valid zero mean. Empty supports are stored as *Q_cm_* = 0, *N_cm_* = 0 and excluded from analyses requiring nonempty measurements. Nonfinite values contribute to neither sum nor count. All cellular measurements use complete supports, including portions outside displayed image crops.

### Directional spatial weighting in HR-IMC

HR-IMC used an optional directional correction based on nuclear elongation and orientation, retaining the same cell centers and candidate sets. For orientation matrix *R_c_* and major-to-minor axis ratio *r_c_*,

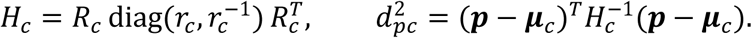

The spatial log weight becomes −(*d_pc_*/*s*)*^α^*, with *r_c_* capped at eight. A cell-specific additive offset matches the total directional weight over its candidate support to the corresponding center-kernel total before intensity scoring. Thus, directional preference changes while total spatial weight is preserved; absolute nuclear size does not enter the kernel.

Nuclear geometry was estimated from component moments after watershed segmentation of summed DNA1 and DNA2 images seeded by cell centers. Morphology and matching criteria restricted the correction to 676 of 4,756 cells; all others retained the center kernel. These auxiliary measurements used no Source-mask boundaries and did not replace the input cell instances.

### Comparison methods

All methods used the same input images and cell instances. Source averaged raw intensities within the supplied masks. Blur and RATI were adapted from spatially smoothed quantification^14^. Blur applied a reflected-edge 3 × 3 box filter before mask averaging. RATI used the median mask mean across ten independent affine perturbations comprising translation, rotation, scaling and shear; cells absent from all transformed samples retained Source values. Transformation settings are specified in the analysis configurations.

The REDSEA-style comparator^8^ used a two-pixel inner boundary band defined by four-connected dilation of background. Contact weights counted background pixels whose 3 × 3 neighborhood contained both cell labels. Each cell’s boundary signal was added to its mask sum, and neighboring boundary signals were subtracted using normalized contact weights. Corrected sums were clipped at zero and divided by the unchanged mask area; isolated cells retained Source values. These comparisons used adapted implementations of the respective approaches.

### Reference-based cohort evaluation

Protein-profile changes were evaluated using 10 predefined marker pairs for Jackson, 10 for Xiao and 14 for Hoch. Each pair linked two nonoverlapping reference groups and one representative marker per group. Within either group, its representative was the expected marker and the paired group’s representative was the other-group marker. Fine annotations supplied both the primary PixAQ anchors and evaluation references; marker-pair definitions were applied only during evaluation.

Within each ROI, each marker was scaled by the 95th percentile of its Source cell means, with that factor fixed across methods. For scaled markers *u_X_*, *u_Y_*, expected-marker share was the equally weighted average of the mean *u_X_*/(*u_X_* + *u_Y_*) in X-reference cells and mean *u_Y_*/(*u_X_* + *u_Y_*) in Y-reference cells. Two measured zero means were assigned a share of 0.5; empty supports were treated as missing.

Each ROI-pair required at least 20 eligible cells in each reference direction, with finite measurements and nonempty supports for both markers under every compared method. Source scales had to be finite and greater than 10^−8^. Methods therefore shared eligible cells while retaining their own pixel supports. Reference pairs, scales and coverage are provided in Supplementary Table 1.

Method-minus-Source share differences were calculated within ROI-pair, summarized by the median across ROIs within patient-pair, and averaged equally across each patient’s eligible pairs. The cohort effect was the median patient effect, expressed in percentage points. Source shares followed the same hierarchy.

Intensity contrasts used direction-specific cell means on the same eligible cells. With a pseudocount of 10^−3^, we calculated log₂ method-to-Source ratios for expected markers and inverse ratios for other-group markers. Reciprocal directions were averaged within ROI-pair and aggregated through the same patient hierarchy. Estimates and intervals were back-transformed to percentage changes, reorienting other-group contrasts and their interval endpoints so that positive values indicated increases for either marker role.

### Spatial support and image analyses

Ownership changes were related to Source boundaries using finite positive raw intensity at pixel-marker positions with different Source and PixAQ owners (Fig. 2d). Distances were measured to the complete Source boundary. Changed intensity was partitioned into transitions between cells, from outside Source masks to a cell, and from a Source cell to an unassigned outcome. ROIs were weighted equally, and the denominator was total intensity at these changed-owner positions.

The within-mask analysis restricted each fine-anchor support to *S_cm_* ∩ *M_c_*, where *M_c_* is the corresponding Source mask (Fig. 2e). Contacting cells had four-connected mask adjacency to a cell in the opposite reference group. Each cell counted once per pair, independently of contact number or its neighbor’s measurement eligibility. Starting from the five-method common eligible set, we required finite, nonempty intersections for both markers; an empty intersection excluded that cell-pair record from all methods. Each ROI-pair retained at least 20 cells per reference direction. Readouts were recalculated on the intersections, while Source used mask means on the same cells. Source scaling and patient aggregation followed the primary benchmark. This restriction changed only the readout domain, without refitting or changing attribution.

Population summaries covered signal sums, pixel counts, means and empty supports (Extended Data Fig. 2). Finite, nonempty Source measurements defined the denominator for empty-support rates. Paired percentage changes additionally required nonempty PixAQ measurements and positive Source signal sums; Source-zero measurements were reported separately. Changes were summarized by medians within ROI-marker and then across ROIs within patient-marker (Supplementary Table 2).

Local image analyses paired signal maps with complete-cell measurements, using shared marker-specific display limits across raw, Source and PixAQ views. Display fields and melanoma cluster profiles were chosen after data inspection. HR-IMC ATP5A/HSP10 fields and cells were chosen using Source signal and spatial position. Cross-marker recipient frequencies included all coordinates with positive raw intensity in both channels within the analyzed field.

### Component and anchor comparisons

For Fig. 3c, we uniformly sampled 20,000 pixel positions from each of the 137 Xiao ROIs and evaluated six markers: CD4, CD8a, CD20, CD68, CD127 and EpCAM. Records required positive raw signal and an alternative candidate from an anchor group different from the nearest cell, yielding 3,928,035 eligible records. The alternative with the highest combined spatial-intensity score was compared with the nearest cell. Pairwise wins required the alternative to outscore the nearest cell; full-competition wins additionally required it to beat every candidate and residual. Rates were averaged equally across the 822 ROI-marker strata. Hexagon colors represent record percentages under the same equal-stratum weighting, with equal weights within each stratum.

Geometry removed candidate-cell intensity scores while retaining spatial weights. Uniform distance replaced each candidate’s spatial weight by the mean over that pixel’s candidates, preserving their total weight. Both controls retained candidate sets and residual competition, and each arm was refitted separately. Shuffled controls permuted fine labels within ROIs, preserving label counts and evaluation references. Common eligibility was applied jointly to Source, fine-anchor Full, Lineage, Fine shuffled, Geometry and Uniform distance. Fine-minus-lineage and fine-minus-shuffled effects were summarized as medians of patient-paired differences.

The matched partition comparison varied only anchor memberships among lineage, annotated cell-type and published cluster partitions. Within each cohort, fits shared images, cells, implementation, numerical settings, initialization procedure, seed and update budget. Source and all three anchor results used common eligible cells, benchmark reference pairs and Source scales (Supplementary Table 3).

### Protein-profile analyses

Pellet profiles were compared within STARLING’s 13 fixed initial clusters. Cluster means for CD20, CD3, CD68 and panCK were divided by each marker’s maximum Source cluster mean, fixed across methods. Dominant-marker margin was the largest minus second-largest scaled mean; a cutoff of 0.15, motivated by STARLING^18^, was used as a descriptive guide for the Source-scaled means in this analysis. Clusters were paired descriptive units from one specimen (Supplementary Table 4).

The melanoma analysis summarized 35 markers across 20 reference groups (Fig. 4e). Each ROI-group-marker required at least 20 cells with finite, nonempty paired measurements. ROI group means were averaged equally, and the PixAQ-minus-Source difference was divided by the population standard deviation of the 20 Source group means for that marker. Additional cluster profiles retained the same 75-cluster partition and common nonempty measurements under Source and both PixAQ anchor arms. Medians of asinh(*Q_cm_*/5) were calculated within ROI-cluster, across ROIs within patient, and then across patients, before division by the marker-specific 99th percentile of Source-cell transformed intensity.

HR-IMC used the PG23 directional fit and standard hard readout. Protein profiles were transformed as asinh(*Q_cm_*/5) and standardized using Source-derived marker means and standard deviations, fixed across methods. Population comparisons retained 2,542 cells with measured Source-mask areas of 500–8,000 pixels in two descriptive groups defined from PG23 Source profiles. These measured areas were distinct from the deposited areas used for anchor construction. ATP5A/HSP10 comparisons paired the same cells and used raw-intensity means over each method’s complete support. Spearman correlations were calculated within ROI and marker, both across the two groups and within each group. Group medians and interquartile ranges were descriptive (Fig. 5d).

### Reference-class prediction and statistical analysis

The auxiliary classifier analysis used 50,755 matched melanoma cells, 35 markers, 137 ROIs and 26 patients to predict five published reference classes (Extended Data Fig. 6a,b). PixAQ was fitted once to the full cohort, and its measurements were held fixed across five patient-held-out classifier folds. Within each fold, cell means were log1p-transformed, the scaler was fitted on training patients, and a class-balanced logistic regression model was trained on the scaled data^34^. Pooled out-of-fold predictions supplied macro-F1, balanced accuracy and per-class recall. Reference classes were obtained by merging Cluster75 through Celltype20 annotations.

Patients were the units for cohort inference. Cohort, ablation, contact-domain and anchor-partition effects used percentile 95% intervals from 10,000 within-cohort patient resamples, preserving method pairing and the specified aggregation. Classifier intervals used 2,000 paired resamples of saved patient confusion matrices without refitting PixAQ or classifiers. ROI-, cluster-, cell- and pixel-level summaries were descriptive. We used nonparametric bootstrap intervals^35^ and performed no null-hypothesis tests.

### Data and code availability

The published datasets analyzed in this study are available from Zenodo: Jackson breast cancer^26^, Xiao melanoma^27^, Hoch melanoma images and masks^28^ and processed data^29^, the STARLING cell-line pellet^30^, and Bollhagen HR-IMC^31^. PixAQ source code will be made publicly available upon journal publication.

## Acknowledgments

This work is supported in part by the Wang Deyao Outstanding Graduate Scholarship Program of Xiamen University (0660-T1062508) and the Xiamen University-Junlong Life collaborative project “Center for Disease Digitalization and Financial Innovation” (20243160C0023).

## Author contributions

Y.Z., X.X., and R.Y. conceived and designed the study. Y.Z. and C.L. developed PixAQ, implemented the computational pipeline, curated the datasets, and performed the computational experiments and downstream analyses. Y.L., G.L., C.Z., Z.H., Z.W., L.Z., and M.T. contributed to methodological discussions, data interpretation, and refinement of the analyses. W.J. provided expertise in mass spectrometry and guidance on experimental data analysis. X.X. and R.Y. supervised the study. Y.Z. drafted the manuscript.

